# Avidity-controlled biotherapeutic delivery systems for the treatment of ischemia-reperfusion acute kidney injury

**DOI:** 10.64898/2026.09.30.755753

**Authors:** Arielle M. D’Elia, Kenneth M. Kim, Alleé Shaw, Sayeh Gholampoor, Jessica Friedrich, Carl R. Russell, Akari J. Seiner, Alexander R. Osidach, Abigail L. McSweeny, Winni Gao, Emma Adolay, Sarah Lynch, Angus Stergar, Evan Rajadhyaksha, Biplab Sarkar, Danielle E. Soranno, Christopher B. Rodell

## Abstract

Regulatory T cells (Tregs) are key mediators of immune resolution and tissue repair following acute kidney injury (AKI), yet strategies to selectively enrich Tregs within the renal microenvironment remain limited. Interleukin-2 (IL-2) is essential for Treg survival and function, but systemic IL-2 delivery is constrained by a short half-life, narrow therapeutic window, and risk of off-target activation of cytotoxic T cells. Effective Treg-based immunomodulation may therefore require both localized cytokine signaling and targeted recruitment of precursor immune populations. Here, we present a sequential, biomaterial-based immunomodulatory strategy that integrates immune cell recruitment with sustained cytokine delivery. An injectable granular hydrogel physically encapsulating the chemokine CCL21 was used to initially recruit CD4+ T cells to the injured kidney. Subsequently, supramolecular guest–host interactions were leveraged to deliver IL-2 modified with the hydrophobic “guest” moiety adamantane, enabling prolonged local cytokine release over 28 days. In healthy mice, we observed early CD4+ T cell recruitment, followed by a pronounced increase in Treg frequency, consistent with immune reprogramming driven by our sequential delivery system. In a murine model of bilateral ischemia–reperfusion (BiAKI), this injectable biomaterial induced a local increase in the proportion of FOXP3-expressing Tregs and a concomitant decrease in the levels of AKI markers. This two-step immunomodulatory approach minimizes acute immune activation while promoting long-term enrichment of Tregs within the renal microenvironment. Together, these findings demonstrate that granular hydrogels with supramolecular host capability can orchestrate immune recruitment and differentiation in a spatiotemporally controlled manner, providing a versatile immunomodulatory platform for applicability in kidney disease.

**HIGHLIGHTS:**

- Acute kidney injury (AKI)-induced inflammation perpetuates renal failure.
- Regulatory T cells (Tregs) are renoprotective post-AKI.
- Granular hydrogels with supramolecular (host) functionality were designed.
- Fast release of CCL21 and slow release of modified IL-2 was achieved.
- Dual-release induced abundant Tregs, protecting against kidney injury.

## 1. INTRODUCTION

Acute kidney injury (AKI) and chronic kidney disease (CKD) are characterized by a robust inflammatory response that contributes to parenchymal damage, impaired repair, and progressive fibrosis[1]. Among immune populations involved in renal injury, CD4^+^ and CD8^+^ effector T cells play a central role in amplifying tissue damage, whereas regulatory T cells (Tregs) mediate immune resolution and tissue repair[2]. Tregs suppress excessive inflammation, promote macrophage polarization toward reparative phenotypes, and support regeneration of tubular epithelial cells[3]. In kidney disease, higher Treg abundance correlates with improved functional outcome, indicating they are an attractive and underexplored therapeutic target for immunomodulatory therapies[4, 5].

Interleukin-2 (IL-2) is indispensable for Treg survival, stability, and function[6, 7]. Low-dose IL-2 therapy has therefore emerged as a promising strategy to selectively expand Tregs without broadly activating effector T cells[8, 9]. However, systemic administration of IL-2 is limited by its short half-life, narrow therapeutic window, and risk of off-target immune activation[10]. These limitations are especially pronounced in solid organs such as the kidneys, where sustained and localized cytokine signaling is difficult to achieve[11, 12]. Thus, there is a critical need for biomaterial-based strategies that not only enhance IL-2 bioavailability but also provide spatio-temporal control over immune modulation.

Beyond cytokine delivery alone, effective Treg-based immunotherapies may require a multistep approach that first recruits appropriate precursor immune populations to injury sites before directing their differentiation and expansion. Chemokines, such as CCL21, play a key role in guiding CD4^+^ T cell migration, making them attractive candidates for immune cell recruitment[13, 14]. Granular hydrogel systems, composed of packed hydrogel microparticles, offer unique advantages for this purpose, including injectability, inherent porosity, and the ability to physically encapsulate and locally retain bioactive factors[15–17]. Such materials can create permissive microenvironments that support immune cell infiltration while maintaining localized chemokine gradients.

Supramolecular biomaterials enable modular and reversible control over protein presentation through noncovalent guest–host interactions[18, 19]. In particular, the interaction between adamantane and cyclodextrin has been widely used to engineer injectable hydrogels with tunable mechanical properties and dynamic cargo retention[20–22]. By modifying therapeutic proteins with guest moieties, supramolecular systems can sustain local signaling without permanent immobilization, preserving protein bioactivity while extending residence time *in vivo*.

Here, we integrate these concepts into a sequential immunomodulatory strategy for treating kidney injury. We employ a granular hydrogel physically encapsulating the chemokine CCL21 to first rapidly recruit CD4^+^ T cells to the injured kidney. Subsequently, we leverage supramolecular guest–host interactions to sustainably deliver IL-2 modified with adamantane, thereby promoting local Treg expansion and differentiation over time. We hypothesized that this two-step approach — immune cell recruitment followed by sustained cytokine signaling — would enable prolonged enrichment of Tregs within the renal microenvironment, preventing the progression of acute kidney injury to chronic kidney disease.

Using a murine model of bilateral warm renal ischemia–reperfusion injury, we examined how our hydrogel modulates the immune response, with a focus on Treg induction. Our findings demonstrate that sustained supramolecular delivery of adamantane-modified IL-2 at an appropriate dose drives a pronounced increase in Treg abundance without inducing cytotoxic T cells. Increased Treg abundance was mirrored by a marked reduction in biomarkers of kidney injury (NGAL, KIM-1), reduced fibrosis, and preservation of kidney function (glomerular filtration rate). These results underscore the importance of local immune modulation and highlight granular, supramolecular hydrogels as a powerful platform for orchestrating immune recruitment, differentiation, and repair in the context of kidney injury.

## 2. MATERIALS AND METHODS

### 2.1 Materials

All reagents and solvents were purchased from Thermo Fisher Scientific unless otherwise indicated.

### 2.2 Granular Hydrogel Fabrication

Granular hydrogels were fabricated based on our previously reported supramolecular hydrogel platform[23]. Briefly, methacrylated dextran (DexMA, 75 kDa) and methacrylated β-cyclodextrin (MeCD, TCI America) were synthesized via esterification with glycidyl methacrylate (GMA, TCI America) and characterized by ^1^H NMR as described. DexMA (0.5 equiv.) and MeCD (0.5 equiv.) were used throughout this study, consistent with the previously reported formulation. Precursor solutions containing 5%_w/v_ DexMA and 10%_w/v_ MeCD with 5 mM lithium phenyl-2,4,6-trimethylbenzoyl phosphinate (LAP, Sigma-Aldrich) were photocrosslinked (10 mW/cm², 5 min) to form bulk hydrogels. Bulk hydrogels were subsequently processed into microgels via extrusion fragmentation (EF) by serial extrusion through progressively smaller gauge needles (18–30G), followed by centrifugation (2000 × *g*, 2 min) three times to induce particle jamming and remove excess supernatant. The resulting microgels exhibited particle sizes and rheological properties consistent with those reported previously[23].

### 2.3 Chemical Synthesis of Adamantane-Modified IL-2 (Ad-IL2)

Recombinant murine interleukin-2 (IL-2, PeproTech 212-12) was modified with adamantane using N-hydroxysuccinimide (NHS) ester chemistry. IL-2 was dissolved in 0.1 M sodium bicarbonate buffer (pH 8.5) at a concentration of 10 mg/mL. Adamantane-PEG-NHS (Ad-PEG-NHS, 5 kDa, RuixiBiotech) was dissolved in anhydrous dimethyl sulfoxide (DMSO, Thomas Scientific) and added at a 5-fold molar excess relative to lysine residues in IL-2. The final DMSO concentration in the reaction mixture did not exceed 10%_v/v_. The reaction was allowed to proceed at room temperature for 4 h under gentle agitation. Following completion of the reaction, 0.1%_w/v_ mouse serum albumin (MSA, Sigma-Aldrich) in Milli-Q water was added to stabilize the modified protein. The reaction mixture was purified by dialysis (3.5–5 kDa MWCO, Spectrum) against Milli-Q water for 48 h, with frequent buffer exchanges. Purified Ad-IL2 was subsequently frozen and lyophilized in low-binding Eppendorf tubes (Snap Cap Low Retention Microcentrifuge Tubes). Lyophilized Ad-IL2 was stored at −80°C until use.

### 2.4 Hydrogel Release Studies

Granular hydrogels (50 µL, n≥3) were loaded with 5 μg of either CCL21, IL-2, or Ad-IL2. Hydrogels were injected into low-binding Eppendorf tubes, and 1 mL of PBS containing 0.5% fatty acid–free BSA (VWR) was added to each tube. Tubes were incubated at 37°C in a humidified incubator with 5% CO_2_. At designated time points (days 1, 2, 4, 7, and twice weekly thereafter until day 28), tubes were briefly centrifuged (1300 × *g*) for 1 min, and the supernatant was collected and replaced. Collected supernatants were stored at −80°C until quantification. Released CCL21 and IL-2 were quantified by ELISA (R&D Systems; MAB4571, M2000), according to the manufacturer’s instructions. Cumulative release was calculated as the sum of protein measured in each collected fraction over time, normalized to total protein recovered.

### 2.5 Animals

All animal procedures were conducted in accordance with the guidelines for the care and use of laboratory animals at Drexel University and Indiana University and were approved by their respective Institutional Animal Care and Use Committees (IACUC Protocol #LA-22-056 at Drexel University; Protocol #22030 at Indiana University). Female C57BL/6J mice (6–8 weeks old) were purchased from Charles River and housed under standard conditions. For kidney studies, we purchased male BALB/c mice (∼8 weeks old) from Jackson Laboratories and housed them under standard conditions. Male mice were selected because female mice are less susceptible to I/R injury[24–26], necessitating prolonged ischemia that would introduce an uncontrolled variable when assessing sex-based effects.

### 2.6 Splenocyte Isolation

Spleens were harvested and immediately placed on ice in RPMI 1640 supplemented with 10% heat-inactivated fetal bovine serum (FBS) and 1% penicillin-streptomycin (pRPMI). Spleens were mechanically homogenized and passed through 70 μm nylon cell strainers. Strainers were rinsed with 5 mL of pRPMI to maximize cell recovery, and cell suspensions centrifuged (300 × *g*, 6 min, 4°C). Supernatants were discarded, and red blood cells lysed by resuspending the cell pellet in 2 mL of ACK lysis buffer for 5 min; this step was repeated as necessary until red blood cells were eliminated. Lysis was quenched with the addition of 8 mL pRPMI, and cells recovered by centrifugation (300 × *g*, 6 min, 4°C). Following removal of the supernatant, cell pellets were resuspended in pRPMI.

### 2.7 Naïve CD4⁺ T Cell Isolation

Naïve CD4⁺ T cells were isolated from splenocytes using magnetic labeling and separation according to the manufacturer’s instructions (MACS, Miltenyi Biotec, Germany). Briefly, splenocyte suspensions were incubated with a biotin-conjugated antibody cocktail targeting non-CD4⁺ T cell populations, followed by anti-biotin magnetic microbeads for negative selection. Labeled cells were passed through MACS separation columns placed in a magnetic field, and the unlabeled naïve CD4⁺ T cells were collected from the flow-through. Isolated cells were washed, counted, and resuspended in RPMI 1640 supplemented with 10% heat-inactivated FBS, 1% penicillin-streptomycin-L-glutamine, 1% MEM nonessential amino acid solution, and 1% sodium pyruvate solution (complete RPMI) for subsequent experiments.

### 2.8 CD4^+^ T Cell Activation and Treg Differentiation

Purified naïve CD4⁺ T cells in complete RPMI were activated overnight with Dynabeads Mouse T-Activator CD3/CD28 at a ratio of 25 μL beads per 1 × 10⁶ T cells, supplemented with 1 μL/mL β-mercaptoethanol (BME) and 2000 U/mL recombinant human IL-2 (carrier-free, BioLegend). For Treg differentiation, activated cells were cultured for 4-5 days at 37°C in a humidified incubator with 5% CO_2_ in the presence of 5 ng/mL recombinant mouse TGF-β1 (carrier-free, BioLegend), with Dynabeads, IL-2, and BME maintained. Media was refreshed halfway through the culture period. At the conclusion of either the overnight CD4⁺ T cell activation or the Treg differentiation culture, Dynabeads were removed using a magnetic separation rack.

To confirm that Ad-IL2 retained bioactivity, Tregs were differentiated in parallel using either unmodified IL-2 or Ad-IL2 under identical conditions, and resulting cells were analyzed by flow cytometry.

### 2.9 Bone Marrow-Derived Macrophage (BMDM) Preparation

BMDMs were obtained from the femurs and tibiae of C57BL/6J female mice as previously described[27]. Briefly, cells were flushed from the bones, filtered through 70 μm strainers, and red blood cells were lysed using ACK buffer. Cells were plated and differentiated for 7 days in Iscove’s Modified Dulbecco’s Medium (IMDM) supplemented with 10% heat-inactivated FBS, 1% penicillin–streptomycin, and 20 ng/mL recombinant murine M-CSF. Differentiated BMDMs were washed and subsequently used in transwell migration assays after 24 h pretreatment with chemokines.

### 2.10 Transwell Migration Assays

Migration of activated CD4⁺ T cells and Tregs was assessed using 5 μm transwell inserts (96-well format, Corning). Prior to the assay, cells were starved for 3 h in chemotaxis media consisting of RPMI 1640 supplemented with 0.5% fatty acid-free BSA (VWR). Inserts were soaked overnight in chemotaxis media. For each assay, 1.25 × 10^5^ cells in 75 μL chemotaxis media were added to the upper chamber, and 235 μL chemotaxis media containing either no stimulus (control), chemokines (IFN-γ, IL-1β, CCL5, CCL21, or CCL22; 100 ng/mL each), or BMDMs pretreated for 24 h with the same chemokines were added to the lower chamber. Plates were incubated for 24 h at 37°C in a humidified incubator with 5% CO_2_. After incubation, cells from both the upper and lower chambers were counted manually using a hemocytometer. Percent migration was calculated as the number of cells in the lower chamber relative to the total number of cells. Cumulative datasets (n = 8) represent independent experiments across biological replicates.

### 2.11 Hydrogel Administration in Healthy Mice

Female mice were subcutaneously injected with 50 µL of granular hydrogel to evaluate dosage and immune response. Three separate studies were performed, each including a control group of mice that did not receive hydrogel injection. To evaluate CCL21 dosage, mice received blank hydrogel, 2.5 µg CCL21, 5 µg CCL21, or 25 µg CCL21 and were sacrificed on day 7 post-injection. To evaluate Ad-IL2 dosage, mice received blank hydrogel, 1 µg Ad-IL2, 5 µg Ad-IL2, or 10 µg Ad-IL2 and were sacrificed on days 3 and 7 post-injection. To evaluate the combination of both, mice received blank hydrogel, 25 µg CCL21, 5 µg Ad-IL2, or [25 µg CCL21 and 5 µg Ad-IL2] combination, and were sacrificed on day 28. At the time of sacrifice, hydrogels, inguinal lymph nodes, and spleens were collected for processing.

### 2.12 Tissue and Hydrogel Processing

Spleens were processed as previously described. Hydrogels and lymph nodes were collected and placed on ice in cold pRPMI. Lymph nodes were mechanically dissociated over pre-scratched 6-well plates before filtration through 70 μm nylon strainers, with red blood cell lysis performed as above. Granular hydrogels were homogenized by repeated pipetting and filtered through stacked 20 μm pluriStrainers (pluriSelect), followed by red blood cell lysis. When necessary, lower filters were pressurized via a 3 mL syringe to facilitate complete filtration. All resulting cell suspensions were used for downstream flow cytometry analyses.

### 2.13 Flow Cytometry

Following tissue and hydrogel processing, cells were counted and resuspended in pRPMI (2 × 10^6^ cells/mL). For intracellular cytokine staining, 2 µL of Cell Activation Cocktail with Brefeldin A (BioLegend) and 1 µL of 1000x monensin solution (BioLegend) were added per mL of cell suspension. Samples were incubated at 37°C for 4 h in a 24-well plate to allow cytokine accumulation. After stimulation, cells were transferred to a 1 mL deep-bottom plate (Azzota Corp) and centrifuged to pellet the cells.

Cells were washed with PBS and stained with LIVE/DEAD Fixable Aqua Dead Cell Stain Kit and TruStain FcX (anti-mouse CD16/32; BioLegend) for 10 min at 4°C. Cells were washed again and incubated with surface antibodies for 30 min at 4°C: Alexa Fluor 700 anti-CD45 (30-F11), APC anti-TCR-β (H57-597), Super Bright 645 anti-CD4 (GK1.5), eFluor 450 anti-CD8a (53-6.7), BV786 anti-CD62L (MEL-14), FITC anti-CD44 (Clone IM7), and PE-Cy7 anti-CD25 (Clone PC61.5). Following surface staining, cells were washed with Flow Cytometry Staining Buffer (eBioscience) and subjected to fixation and permeabilization (overnight, 4°C) using the FOXP3/Transcription Factor Staining Buffer Set (eBioscience). After two washes with permeabilization buffer, cells were incubated with intracellular antibodies for 1 h at 4°C: PE-eFluor 610 anti-FOXP3 (FJK-16s) and PE anti-Granzyme B (QA16A02). Cells were washed once with permeabilization buffer, resuspended in Flow Cytometry Staining Buffer, and analyzed on a BD FACSymphony™ A1 flow cytometer. Single-stained compensation controls were prepared using compensation beads and used to calculate compensation matrices prior to acquisition. Data were analyzed using FlowJo software (version 10).

For the CCL21 dosage and Ad-IL2 bioactivity studies, intracellular cytokine staining was not performed, and Granzyme B was not assessed. In these experiments, CD62L was stained using PE anti-CD62L (MEL-14) in place of BV786.

### 2.14 Renal Hydrogel Injection in Healthy Mice

Healthy male mice received a bilateral subcapsular injection of either 15 µL hydrogel or saline. Kidney function was monitored at baseline, day 3, and day 28 post-injection using transcutaneous glomerular filtration rate (tGFR) measurements. At the time of sacrifice, serum was collected for measurement of blood urea nitrogen (BUN), creatinine, and cystatin C levels. Harvested kidneys were sectioned and stained with picrosirius red (PSR). Fibrotic regions were quantified using Celleste software to determine the fibrosis area, expressed as a percentage of whole tissue area examined.

### 2.15 Hydrogel Delivery in AKI Mice

Male mice were subjected to warm bilateral ischemia– reperfusion acute kidney injury (AKI). After surgical dissection via a dorsal approach, both the left and right renal pedicles were clamped for 25 minutes to restrict blood flow and then unclamped. Reperfusion was confirmed visually prior to surgical closure. Three days post-injury, mice received bilateral 15 µL subcapsular renal injections containing either saline, unloaded hydrogel, hydrogel loaded with CCL21, hydrogel loaded with Ad-IL2, or a combination hydrogel containing both CCL21 and Ad-IL2.

tGFR was measured at baseline (pre-injury), day 4 post-injury, and prior to euthanasia on day 28. Serial blood samples were collected via retro-orbital bleeding to measure blood urea nitrogen (BUN). Urine was also collected for measurement of KIM-1 and NGAL via ELISA (R&D Systems; MKM100 and MLCN20). At the time of euthanasia, kidneys were bisected, with one half stored in ethanol and the other half flash-frozen for downstream analyses.

### 2.16 Histology

Kidney sections were stained with PSR and imaged at 10x magnification using an ECHO Revolution microscope (ECHO, San Diego, CA, USA). Birefringence on PSR-stained slides was captured using a circularly polarizing light condenser attachment, isolating fibrillar collagen fibers. Quantification was performed using ImageJ (version 1.54). For each section, the outer cortical rim (∼4–5 glomeruli depth) was isolated and the percent birefringent area was thresholded and calculated within that region.

For immunofluorescence studies, kidneys were fixed, paraffin-embedded, and sectioned at 5 μm thickness. Sections were deparaffinized in xylene and rehydrated through a graded ethanol series. Antigen retrieval was performed in citrate buffer (10 mM, pH 6.0) using a pressure cooker for 8–10 min. After cooling to room temperature, sections were washed in PBS and blocked with bovine serum albumin (BSA) in PBS to reduce nonspecific binding. Sections were then incubated overnight at 4°C with primary antibodies diluted in blocking buffer. Following washes with PBS, sections were incubated for 1 h at room temperature with fluorophore-conjugated secondary antibodies, washed again, and counterstained with DAPI to label nuclei. Finally, slides were mounted with mounting medium and imaged using a fluorescence microscope.

Quantification of immunofluorescence images was performed using QuPath-0.7.0. Fluorescent images were thresholded to define positive signal, and positively stained cells were identified and quantified. For each section, ten circular regions of interest (ROIs) were drawn within the renal cortex, and positive cells were quantified within these ROIs.

### 2.17 Statistical Analysis

Data are presented as mean ± standard deviation or standard error of the mean, as indicated in figure legends. Statistical analyses were performed with GraphPad Prism v11.0.0 using analysis of variance (ANOVA) where appropriate with *post-hoc* Tukey’s honest significant difference (HSD) test. For Ad-IL2 release studies, a two-way ANOVA was performed, followed by Bonferroni’s multiple comparisons *post-hoc* test. For studies involving unloaded hydrogel injection into healthy mouse kidneys, a two-way ANOVA was performed followed by Fisher’s least significant difference (LSD) *post-hoc* test. Normality was confirmed by Shapiro–Wilk test.

## 3. RESULTS

### 3.1 Design of a supramolecular hydrogel platform for T cell recruitment and Treg induction

We engineered a granular hydrogel depot designed to provide temporally controlled immunomodulatory cues for local Treg enrichment in the context of kidney injury (**Fig. 1**). The system was designed to first release a chemokine to recruit CD4⁺ T cells to the site of injury, followed by sustained delivery of IL-2 to promote differentiation and expansion of Tregs. This sequential strategy aims to locally enrich Tregs that suppress inflammation to limit progression from AKI to CKD.

**Figure 1.**
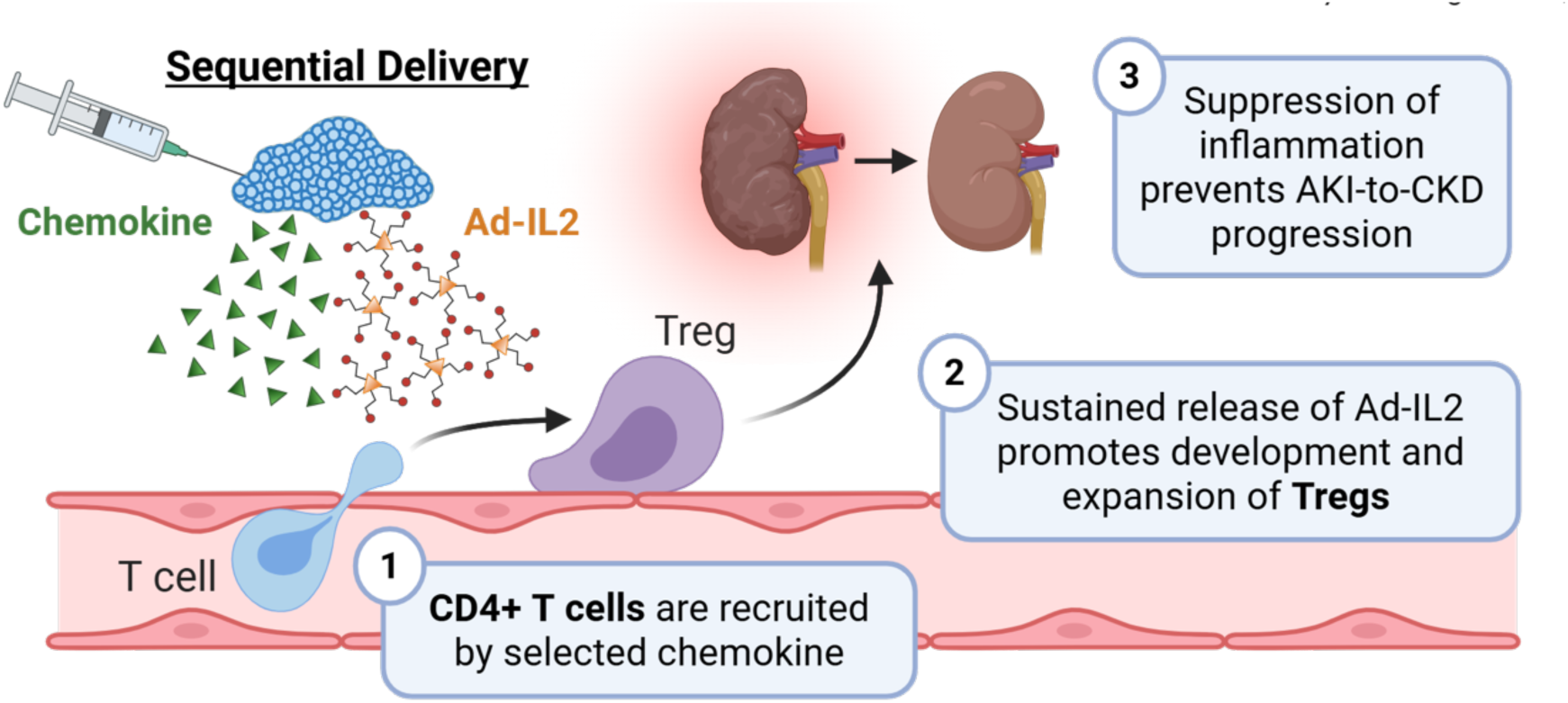
Schematic of the hydrogel-based local delivery system for Treg induction in kidney injury. A granular hydrogel depot is engineered to release therapeutic cues with temporal control. In the initial phase, the hydrogel delivers a chemokine to recruit CD4⁺ T cells to the site of ischemic kidney injury. Subsequently, the material provides sustained release of IL-2 to drive differentiation and expansion of regulatory T cells (Tregs). The resulting enrichment of Tregs at the site of injury suppresses inflammation and is designed to limit progression from acute kidney injury (AKI) to chronic kidney disease (CKD).

### 3.2 CCL21 release promotes CD4⁺ T cell recruitment *in vitro* and *in vivo*

Naïve CD4⁺ T cells were isolated from murine spleens via negative selection and subjected to activation or Treg induction protocols (**Fig. 2A**). To identify a chemokine capable of recruiting CD4⁺ T cells and Tregs, we evaluated several candidate lymphocyte-chemotactic signals: IFN-γ, IL-1β, CCL5, CCL21, and CCL22. Transwell migration assays revealed that CCL21 induced significantly greater migration of both CD4⁺ T cells and Tregs compared with the other chemokines tested (**Fig. 2C-D**), with CD4⁺ T cell migration increasing more than 3-fold and Treg migration increasing more than 2-fold compared to other chemokines. Similar trends were observed when BMDMs were included as intermediate chemoattractant sources **(Fig. S1)**. Based on these results, CCL21 was selected for incorporation into the hydrogel delivery system.

**Figure 2.**
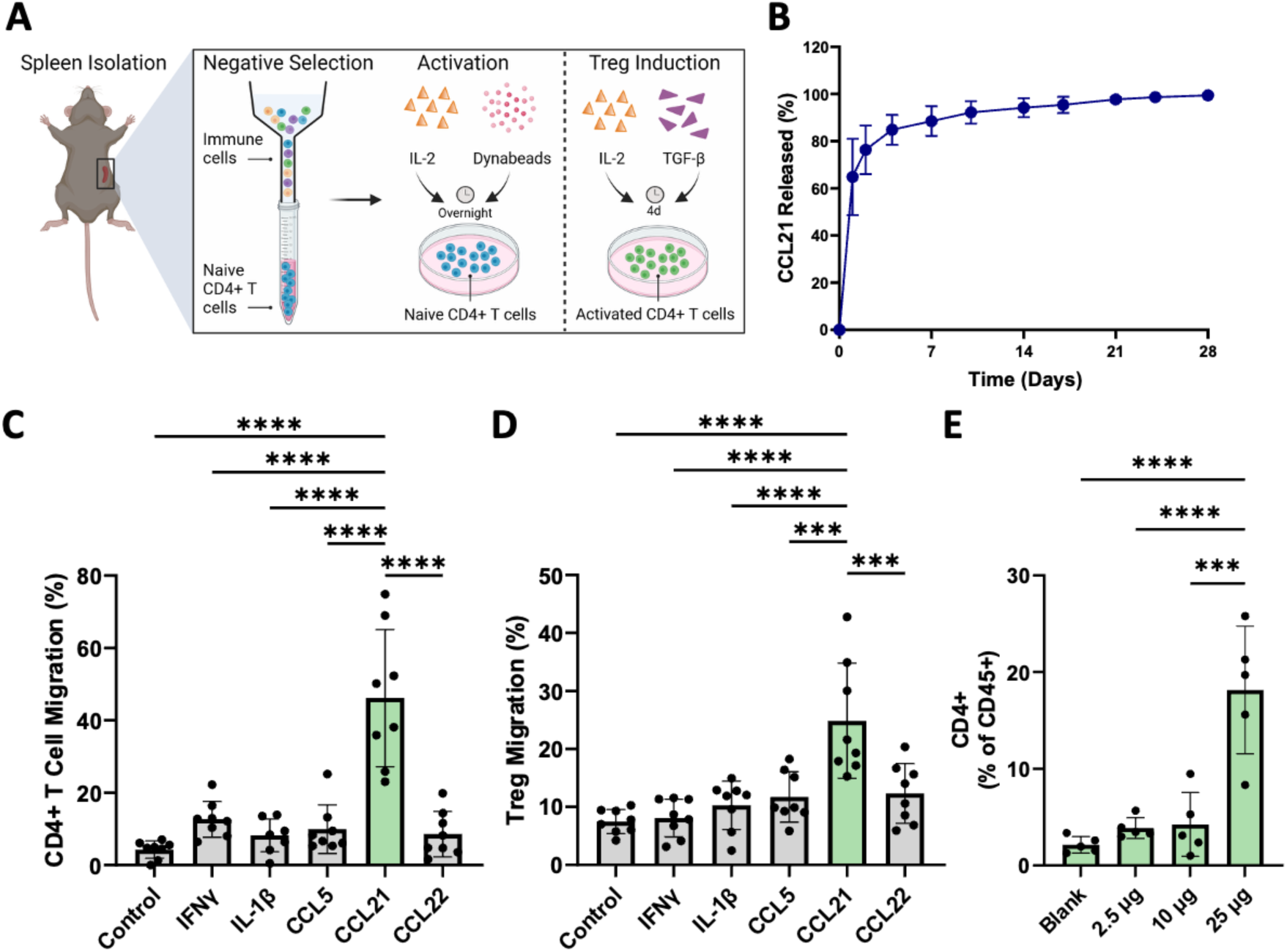
Isolation, migration, and T cell presence. **(A)** Schematic of spleen isolation and naïve CD4+ T cell purification via negative selection, followed by overnight activation or Treg induction. **(B)** *In vitro* cumulative percentage of CCL21 released from granular hydrogels over 28 days; values represent mean ± SD, n=4. Transwell migration assays of **(C)** CD4+ T cells and **(D)** Tregs toward chemoattractants. Data are pooled from two independent experiments (n = 4 per experiment; total n = 8); ***p<0.001, ****p<0.0001; one-way ANOVA, Tukey’s HSD. **(E)** CD4+ T cell population from hydrogel explants on day 7 after *in vivo* injection. Data represent mean ± SD, n=5; ***p<0.001, ****p<0.0001; one-way ANOVA, Tukey’s HSD.

To evaluate chemokine delivery from the hydrogel, release of CCL21 from granular hydrogels was measured *in vitro*. The majority of CCL21 was released within the first week, with approximately 88% released by day 7 (**Fig. 2B**). To determine whether recruitment occurred *in vivo*, hydrogels were explanted 7 days after subcutaneous injection above the kidneys in healthy mice. A significant increase in the percentage of CD4⁺ T cells among recovered immune cells was observed in hydrogels containing 25 µg CCL21 compared to those containing lower doses, reaching approximately 18% of all CD45^+^ immune cells (**Fig. 2E**). Together, these data demonstrate that CCL21 release from the granular hydrogel promotes CD4⁺ T cell recruitment both *in vitro* and *in vivo*.

### 3.3 Adamantane modification enables sustained IL-2 release without impairing Treg induction

To enable sustained cytokine delivery from the hydrogel platform, IL-2 was modified with adamantane (Ad-IL2) via NHS chemistry (**Fig. 3A**), allowing avidity-driven supramolecular interactions within the hydrogel network. To evaluate whether chemical modification affected IL-2 activity, CD4⁺ T cells were induced toward a Treg phenotype in the presence of unmodified IL-2 or Ad-IL2. Ad-IL2 promoted dose-dependent increases in the percentage of Tregs among TCR-β⁺ cells, indicating that IL-2 retained biological activity following Ad conjugation (**Fig. 3B**). *In vitro* release studies comparing IL-2 and Ad-IL2 release from granular hydrogels demonstrated significantly prolonged retention of Ad-IL2 within the hydrogel compared to unmodified IL-2 over 28 days (**Fig. 3C**). Unmodified IL-2 exhibited rapid release, with more than 90% released within the first day. In contrast, Ad-IL2 demonstrated sustained release over several weeks, consistent with supramolecular avidity-mediated release.

**Figure 3.**
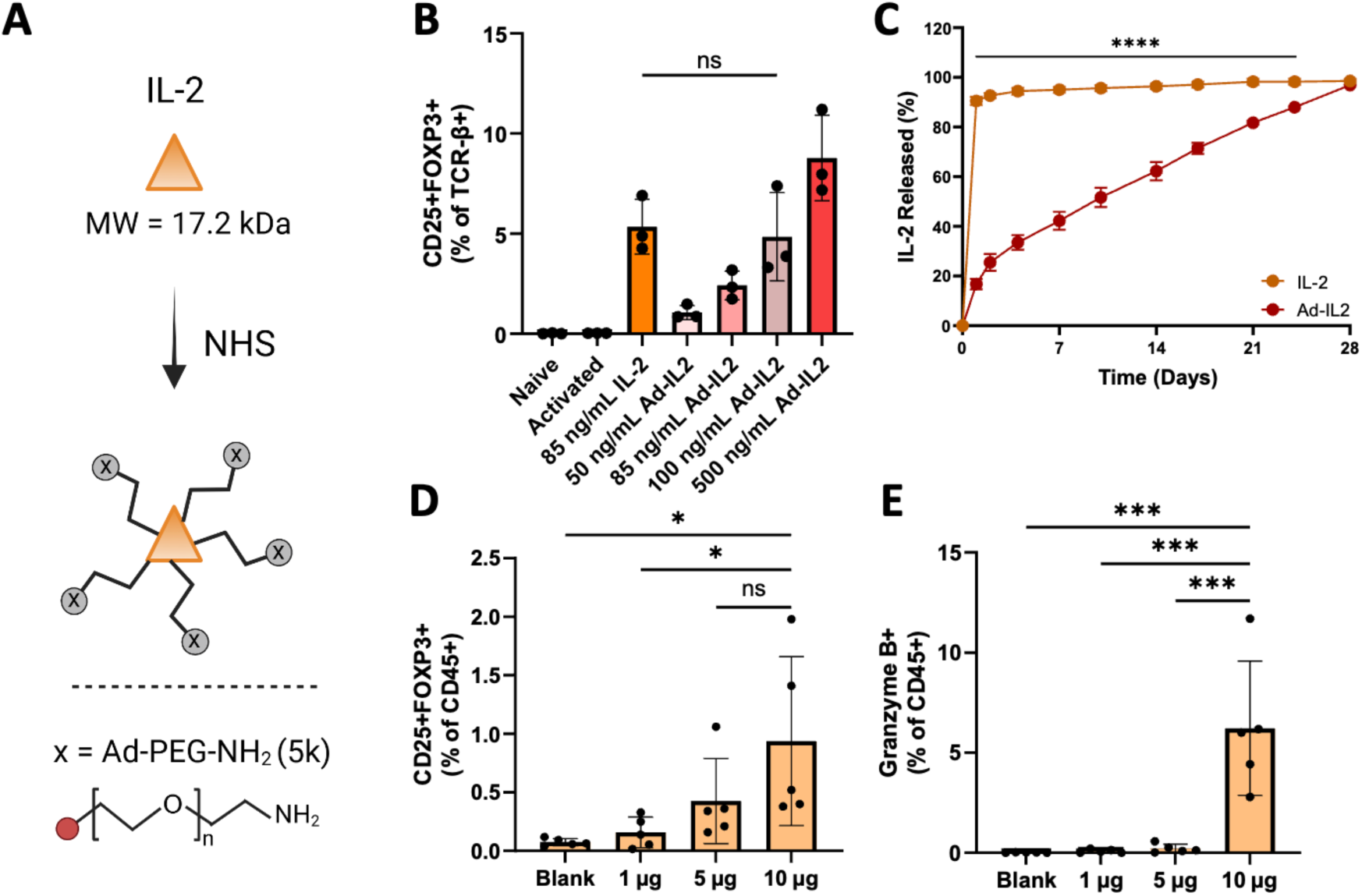
Adamantane modification of IL-2 and its effects on T cell induction. **(A)** Schematic of adamantane attachment to IL-2 via NHS chemistry. **(B)** Percentage of Tregs among TCR-β+ cells following induction with Ad-IL2 *in vitro*; ns, not significant. **(C)** *In vitro* cumulative release of IL-2 and Ad-IL2 released from granular hydrogels over 28 days; values represent mean ± SD; n ≥ 3; ****p<0.0001; RM two-way ANOVA, Bonferroni correction. Percentage of **(D)** Tregs and **(E)** granzyme B+ T cells among all CD45+ immune cells within hydrogels explanted from healthy mice on day 7 after injection with 1, 5, or 10 µg Ad-IL2. Data represent mean ± SD, n=5; *p<0.05, ***p<0.001; one-way ANOVA, Tukey’s HSD.

To determine an appropriate *in vivo* dose, hydrogels containing 1, 5, or 10 µg Ad-IL2 were administered subcutaneously in healthy mice. Increasing Ad-IL2 dosage elevated the percentage of Tregs among CD45⁺ immune cells within hydrogel explants at day 7 (**Fig. 3D**). However, increasing Ad-IL2 dose also increased granzyme B⁺ T cells among CD45⁺ immune cells, with 10 µg producing significantly greater abundance of cytotoxic T cells (more than 30-fold higher than that in lower doses) (**Fig. 3E**). Consistent with these findings, systemic immune compartments also exhibited increased cytotoxic activation at the highest dose, with 10 µg Ad-IL2 significantly increasing the percentage of granzyme B⁺ T cells in both the inguinal lymph nodes and spleen (**Fig. S2A–B**). Based on these results, 5 µg Ad-IL2 was selected for subsequent studies.

### 3.4 Combination delivery of CCL21 and Ad-IL2 enhances local Treg enrichment without promoting cytotoxic T cell activation

The granular hydrogel is designed for temporally controlled cytokine release, with fast-releasing CCL21 and slow-releasing Ad-IL2 through supramolecular adamantane–cyclodextrin interactions (**Fig. 4A**). To assess immune recruitment and programming, CD4⁺ T cells and Tregs were quantified in hydrogel explants over 28 days. Only hydrogels containing CCL21 increased CD4⁺ T cells on day 7, confirming that CCL21 rapidly recruits precursor T cells that can later differentiate into Tregs (**Fig. S6A**). By day 28, the combination hydrogel maintained an elevated proportion of CD4⁺ T cells (**Fig. S6B**), indicating persistent recruitment or retention of helper T cells in the local microenvironment.

**Figure 4.**
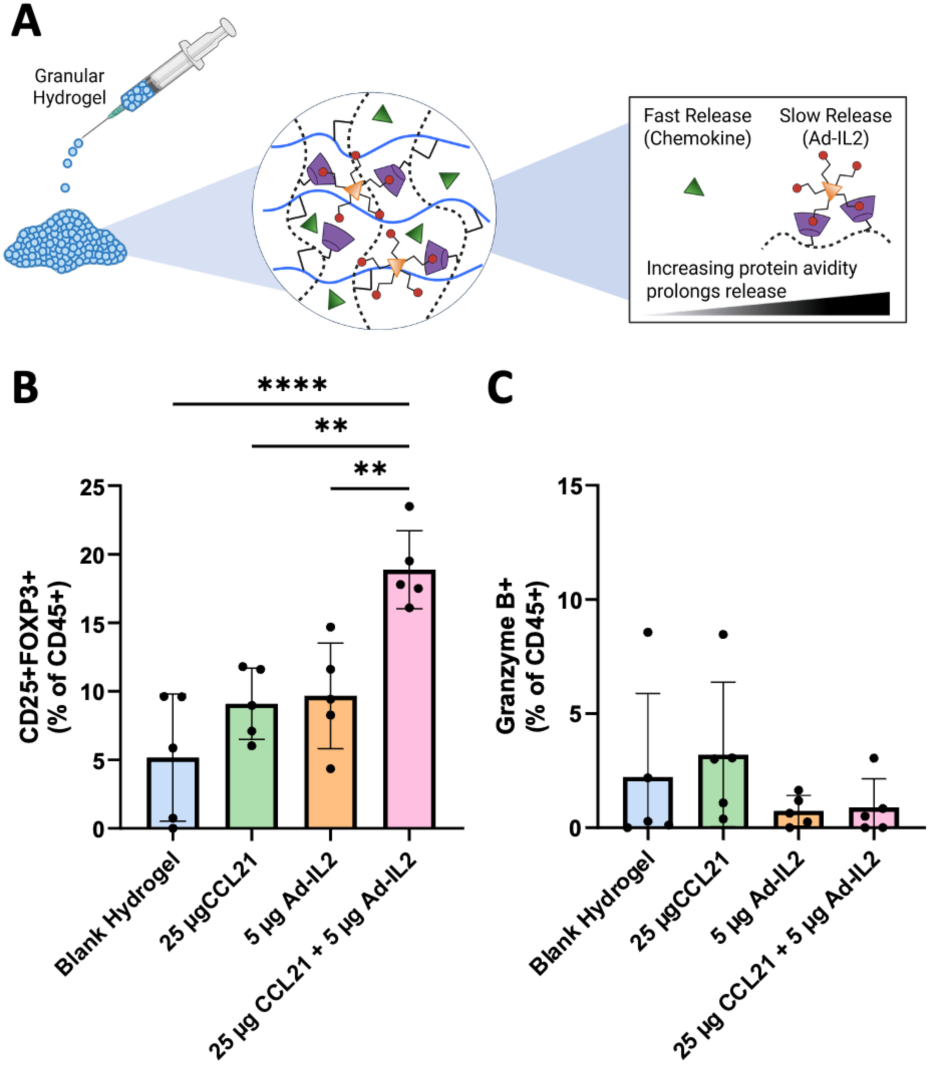
Immune cell analysis of CCL21 and Ad-IL2 loaded hydrogels *in vivo*. **(A)** Overview of the granular hydrogel system and its underlying chemical interactions. The chemokine is rapidly released to promote early T cell recruitment, whereas Ad-IL2 exhibits sustained release mediated by avidity-driven supramolecular interactions. Percentage of **(B)** Tregs and **(C)** granzyme B+ T cells among CD45+ immune cells in blank, 25 µg CCL21, 5 µg Ad-IL2, and combination hydrogels on day 28. Data represent mean ± SD, n=5; **p<0.01, ****p<0.0001; one-way ANOVA, Tukey’s HSD.

Consistent with T cell recruitment, the combination of 25 µg CCL21 and 5 µg Ad-IL2 significantly increased Treg percentages among CD45⁺ immune cells in the hydrogel at day 28 to approximately 18%, outperforming hydrogels delivering either factor alone (**Fig. 4B**). Granzyme B⁺ cytotoxic T cell levels remained very low (fewer than 1% of CD45⁺ immune cells) in all Ad-IL2-containing groups at day 28 (**Fig. 4C**), demonstrating selective expansion of regulatory cells without induction of cytotoxic responses. Systemic analyses further confirmed the localized observations. In the spleen, the combination hydrogel increased Treg percentages on day 28 (**Fig. S6C**). In lymph nodes, inclusion of Ad-IL2 significantly increased Tregs on day 28 (**Fig. S6D**), supporting the idea that slow-release Ad-IL2 sustains regulatory T cell programming beyond the local hydrogel site.

### 3.5 Granular hydrogel delivery to the kidney does not impair renal function

To evaluate the safety of hydrogel delivery to the kidney, granular hydrogels were administered via subcapsular injection in healthy mice, followed by longitudinal assessment of glomerular filtration rate (GFR). Following hydrogel administration, transcutaneous glomerular filtration rate (tGFR) was measured, and serum and kidney tissue were collected for biomarker assessment and histological analysis at day 28 (**Fig. 5A**). Histological evaluation using PSR staining revealed no observable differences in collagen deposition between saline-injected and hydrogel-injected kidneys, indicating that hydrogel administration did not induce fibrosis within the renal tissue (**Fig. 5B**).

**Figure 5.**
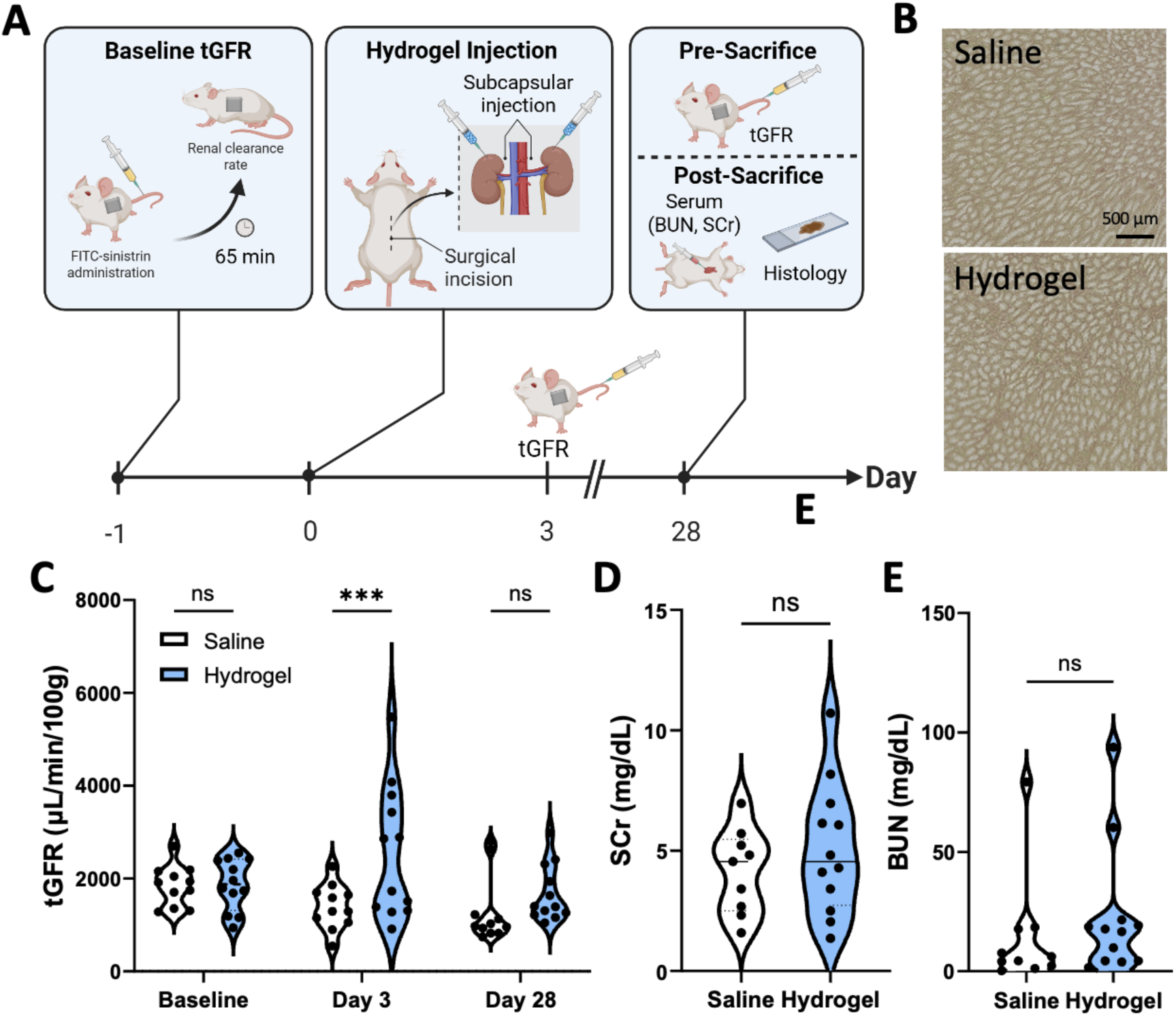
Biocompatibility of granular hydrogel following subcapsular kidney injection in mice. **(A)** Schematic of granular hydrogel delivery via subcapsular injection into healthy mouse kidneys. **(B)** Representative Picrosirius Red (PSR) staining of kidney sections after saline or hydrogel injection. Measurements of **(C)** transcutaneous glomerular filtration rate (tGFR), (**D)** serum creatinine (SCr), and **(E)** blood urea nitrogen (BUN) in saline and hydrogel injected mice. Data represent mean ± SD, n ≥ 11; ***p<0.001; ns, not significant; two-way ANOVA, Fisher’s LSD test.

tGFR measurements revealed a transient increase on day 3 following hydrogel injection compared to saline controls, suggesting an early compensatory response to the material. By day 28, tGFR values were comparable between groups, indicating full recovery and preservation of GFR (**Fig. 5C**). Consistent with these findings, serum creatinine (SCr) and blood urea nitrogen (BUN) showed no significant differences between saline- and hydrogel-treated mice at day 28 post-injection (**Fig. 5D-E**). Together, these data demonstrate that subcapsular administration of the granular hydrogel does not induce fibrotic remodeling nor impair renal function in healthy kidneys.

### 3.6 Supramolecular hydrogel treatment preserves renal function following ischemia– reperfusion (I/R) injury

To evaluate the therapeutic potential of the granular hydrogels to attenuate AKI-to-CKD progression, mice underwent bilateral renal I/R injury (BiAKI), followed by hydrogel administration on day 3 and subsequent assessment at day 28 (**Fig. 6A**). Urinary neutrophil gelatinase-associated lipocalin (NGAL), a marker of kidney injury, was significantly reduced in the combination hydrogel group compared to blank controls, with a >4.5-fold decrease observed (**Fig. 6B**). Urinary kidney injury molecule-1 (KIM-1) levels did not differ significantly between groups (**Fig. S7A**). Renal function assessment revealed that tGFR was significantly higher in mice treated with the combination hydrogel compared to blank hydrogel controls (**Fig. 6C**). Histological evaluation confirmed that hydrogel administration did not induce fibrosis. PSR staining of kidney sections under polarized light showed no significant differences in percent birefringent area within the outer cortical rim between hydrogel-treated and control kidneys (**Fig. S7B**), indicating that the hydrogel does not promote collagen deposition or fibrotic remodeling.

**Figure 6.**
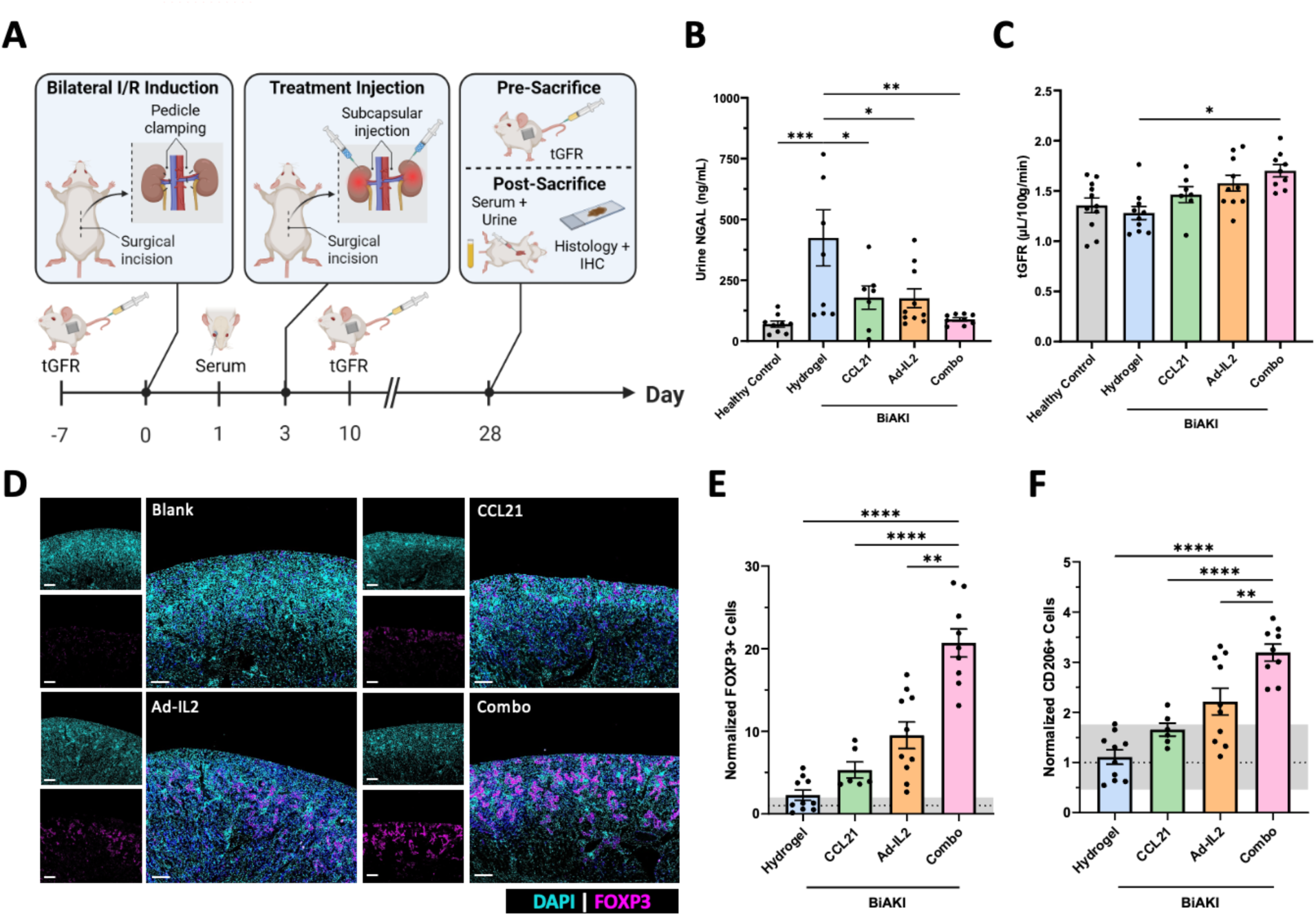
Granular hydrogel treatment preserves renal function following bilateral ischemia–reperfusion injury (BiAKI). **(A**) Schematic of the bilateral renal ischemia–reperfusion (I/R) acute kidney injury (AKI) model. **(B)** Urinary neutrophil gelatinase-associated lipocalin (NGAL) and **(C)** transcutaneous glomerular filtration rate (tGFR) levels at day 28 post-BiAKI induction. **(D)** Representative immunofluorescence images of BiAKI kidneys treated with blank, CCL21, Ad-IL2, or combination hydrogel. Images show merged DAPI and FOXP3 staining from the renal cortex, with corresponding single-channel views (DAPI and FOXP3) from the same field of view. Scale bar 250 µm. Quantification of normalized **(E)** FOXP3⁺ and **(F)** CD206⁺ cells from immunofluorescence analysis. Data represent mean ± SEM, n ≥ 6; *p<0.05, **p<0.01, ***p<0.001, ****p<0.0001; one-way ANOVA, Tukey’s HSD.

Assessment of the local immune response revealed that the combination hydrogel treatment markedly increased renal Treg populations following BiAKI. Whole kidney immunofluorescence imaging demonstrated minimal immune cell presence in the medulla, with immune cells predominantly localized to the cortex (**Fig. S8**). Upon examination of the renal cortex, a pronounced increase in FOXP3⁺ cell presence was observed in the combination hydrogel group compared to controls (**Fig. 6D**). Quantification confirmed a significant increase in Tregs, rising from 0.18% in the blank hydrogel group to 1.56% in the combination hydrogel group, corresponding to a 20-fold increase (**Fig. 6E**).

In parallel, macrophage polarization was assessed within the injured kidney (**Fig. S9**). Anti-inflammatory CD206⁺ macrophages were significantly increased from 7.69% in the blank hydrogel group to 22.81% in the combination hydrogel group, representing a 3-fold increase (**Fig. 6F**); however, the relative abundance of pro-inflammatory iNOS+ macrophages was minimally affected at these later timepoints (**Fig. S7C**). Together, these findings demonstrate that the combination hydrogel improves renal function and injury markers after BiAKI while promoting an immunoregulatory microenvironment characterized by enhanced Treg recruitment and a shift in macrophage polarization toward a reparative phenotype.

## 4. Discussion

The granular hydrogel platform was designed to deliver immune signals in a temporally coordinated manner, first recruiting CD4⁺ T cells and subsequently promoting their differentiation into Tregs through sustained Ad-IL2 release. This strategy enables the local enrichment of regulatory immune populations within the injured kidney microenvironment, concentrating immunomodulatory signals at the site of injury while minimizing systemic exposure and off-target effects. Since Tregs play a critical role in suppressing inflammation and limiting fibrosis following kidney injury[28], enhancing their local abundance represents a promising approach to prevent the progression from AKI to CKD.

Local chemokine delivery allows for the establishment of spatially confined gradients, which are critical for directional leukocyte migration. The robust recruitment observed with CCL21 aligns with its known role in CCR7-mediated T cell trafficking and lymphoid homing[13]. CCR7 signaling regulates lymphocyte migration along chemokine gradients, and because Tregs also express CCR7, localized CCL21 release effectively recruits both CD4⁺ T cell precursors and Tregs to the hydrogel depot. Early, rapid chemokine release is advantageous, as initial immune cell recruitment after injury helps establish a regulatory microenvironment prior to sustained cytokine signaling. By locally expanding the pool of CD4⁺ precursor cells, CCL21 delivery enhances the efficiency of subsequent IL-2–mediated Treg differentiation.

Localized cytokine delivery has been shown to modulate renal inflammation and improve tissue repair, for example through IL-10 delivery[29–31]. Many hydrogel-based cytokine delivery platforms rely on pre-loaded factors, where efficacy is dictated by initial loading and passive release kinetics, which can result in rapid payload depletion and diminished long-term biological effect[32, 33]. Other platforms incorporate supramolecular or affinity-based interactions to prolong release[34], microgel–hydrogel composites to decouple injectability from therapeutic release[35], or stimuli-responsive mechanisms that trigger on-demand delivery[36–38]. While these strategies enhance release control, they are limited by either invasiveness, mechanical complexity, or lack of active recruitment of immune cells.

In contrast, our hydrogel is designed to retain IL-2 through direct supramolecular interactions, where guest–host binding introduces avidity arising from multivalent interactions between IL-2– conjugated adamantane groups and cyclodextrin moieties within the hydrogel network, thereby enabling tunable cytokine sequestration and release kinetics. This multivalent binding behavior increases the effective local residence time of IL-2 within the hydrogel network, allowing for sustained, bioactive presentation that continuously signals to recruited T cells. This design promotes ongoing Treg induction without requiring repeated administration or invasive implantation. The hydrogel’s injectable, minimally invasive format allows conformal delivery to complex tissue geometries while maintaining tunable release kinetics, combining active immune cell recruitment, sustained cytokine presentation, and *in situ* immune programming into a single platform.

Low-dose IL-2 therapy preferentially expands Tregs due to their high-affinity IL-2 receptor[39], but systemic administration is constrained by rapid clearance and potential off-target effects. Chemical modification of IL-2 with adamantane yielded Ad-IL2 that forms supramolecular guest– host interactions with the cyclodextrin moieties in the granular hydrogel, dramatically prolonging local retention compared to unmodified IL-2. Importantly, Ad-IL2 retains biological activity and supports dose-dependent Treg induction. Our data indicate that sustained Ad-IL2 presentation amplifies Treg abundance over several weeks; however, careful dosing is critical to avoid excessive activation of cytotoxic T cells, as higher IL-2 levels have been shown to increase granzyme B⁺ T cells in addition to expanding Tregs[40], underscoring the importance of balancing regulatory and effector immune signaling.

The combination of fast-releasing CCL21 and slowly released Ad-IL2 was synergistic, producing higher and more sustained Treg percentages than either factor alone. Early CD4⁺ T cell recruitment establishes a robust precursor pool, while continued presence of these helper T cells may facilitate ongoing Treg-mediated immune regulation and localized tissue repair. Importantly, Treg enrichment did not coincide with increased cytotoxic T cell activity, demonstrating selective enhancement of regulatory immune responses. Trends toward increased Tregs in secondary lymphoid organs suggest potential systemic benefits, such as mitigating inflammatory spillover and preventing secondary tissue damage. Future studies examining longer-term immune responses, repeated dosing, and additional time points will further clarify potential systemic immune effects and help define immune dynamics during early and later stages of treatment, supporting the controlled, spatiotemporal cytokine release enabled by the hydrogel platform.

The safety of biomaterial delivery to the kidney is a critical consideration for translational applications. Subcapsular injection of the granular hydrogel was well tolerated in healthy kidneys, with no evidence of fibrosis or increased collagen deposition, indicating that the material does not induce fibrotic remodeling within the renal microenvironment. Renal function was preserved over 28 days, as assessed by tGFR, serum creatinine, and BUN, supporting the biocompatibility of the hydrogel and its translational potential. Although a transient reduction in tGFR was observed shortly after injection, glomerular filtration capacity recovered by the study endpoint, suggesting this early change reflects a temporary physiological response to hydrogel placement rather than sustained injury. The absence of persistent alterations in standard clinical biomarkers further confirms that the hydrogel does not impair kidney function.

In the context of AKI, the combination hydrogel preserved renal function and reduced markers of kidney injury. Increased tGFR reflects enhanced glomerular filtration, while reductions in urinary NGAL and trends toward lower KIM-1 levels indicate decreased tubular injury and epithelial stress. The absence of fibrosis and maintenance of renal biomarkers underscore the platform’s safety, highlighting its suitability for translational kidney therapies that require local immune modulation to provide renoprotective effects without compromising organ function.

Beyond functional recovery, the combination hydrogel induced a pronounced immunomodulatory response characterized by increased renal FOXP3⁺ Treg infiltration. This local enrichment of Tregs is particularly significant given their established role in suppressing inflammation, limiting immune-mediated tissue damage, and promoting tissue repair in injured organs. Consistent with this, macrophage polarization within the kidney was shifted toward a reparative phenotype, with a moderate increase in CD206⁺ macrophages observed in the combination hydrogel group. Although iNOS expression was not significantly altered, the coordinated increase in Tregs and CD206⁺ macrophages suggests the establishment of an immunoregulatory microenvironment. These findings are consistent with emerging evidence that local Treg delivery can drive tissue repair and modulate innate immune cell behavior, including macrophage polarization, in injured tissues[41]. However, the driving signals that may coordinate T cell and macrophage response were not examined here. Together, these results support the concept that the granular hydrogel enables a localized Treg–macrophage immunoregulatory axis, in which Tregs may contribute to the recruitment or polarization of reparative macrophages, thereby enhancing renal recovery following injury. By enabling targeted immune modulation while preserving renal function and structure, this system provides a minimally invasive platform for harnessing endogenous repair mechanisms in kidney disease.

Important towards both the interpretation of results and translational relevance is the timing of administration relative to the initial injury. In the present study, the hydrogel was administered immediately following ischemia–reperfusion injury. This timing was selected to intervene during the period in which immune cell recruitment, cytokine production, and tissue damage are actively developing. By establishing a local source of CCL21 and IL-2 at the time of injury, the therapeutic strategy aimed to influence the subsequent immune response before maladaptive chronic inflammation and fibrosis are established. While this approach demonstrated therapeutic benefit as a critical proof of principle, future studies should investigate delayed administration time points to better reflect clinical scenarios. Towards this point, numerous therapeutics have shown promising results in preclinical studies, but attempts to translate such therapies to the clinic have failed[42]. To overcome translational barriers, the NIDDK convened the “AKI Outcomes: Overcoming Barriers in AKI” workshop in 2015[43]. Key aspects of these expert recommendations are implemented in our experimental approach, including: 1) use of a validated, clinically relevant animal model (bilateral ischemia–reperfusion AKI), 2) measurement of glomerular filtration rate rather than reliance upon estimates of kidney function via biomarkers (tGFR measurement), and 3) considering feasibility of therapeutic intervention (e.g., minimally-invasive injection in a procedure similar to percutaneous renal biopsy, which can be performed in children and adults under conscious sedation). Notably, they also recommend matching the preclinical intervention to a clinically feasible and relevant time-point for human intervention; a day 3 time-point would be most pertinent for hydrogel delivery, as it is a clinically relevant time-point for therapeutic intervention; most AKI episodes are diagnosed 48–72 hours after injury. The use of injectable hydrogels for T cell recruitment and subsequent Treg induction represents a novel, yet practical approach to treat AKI and prevent CKD.

Collectively, these findings demonstrate that spatiotemporal control of multiple immunomodulatory factors within a single, injectable hydrogel can program immune cell populations more effectively than single-factor or passive-release approaches. By integrating rapid precursor cell recruitment, sustained cytokine signaling, and selective Treg expansion, this platform offers a mechanistically and translationally distinct strategy for localized immunomodulation in the context of kidney injury. Its minimally invasive, tunable, and modular design provides a foundation for therapeutic applications that require prolonged regulatory immune activity while minimizing off-target cytotoxicity, positioning this hydrogel as a promising platform for treating AKI, preventing CKD progression, and potentially addressing maladaptive tissue remodeling in other pro-inflammatory disorders.

## 5. Conclusion

We have developed an injectable granular hydrogel platform capable of temporally coordinated immunomodulation for kidney injury treatment. By rapidly recruiting CD4⁺ T cells through CCL21 and sustaining local Treg differentiation via supramolecularly retained Ad-IL2, the hydrogel achieves selective expansion of regulatory immune populations while minimizing cytotoxic responses. This dual-action strategy also promotes a pro-repair macrophage phenotype, reduces inflammatory injury markers, and preserves renal function without inducing fibrosis, demonstrating both efficacy and biocompatibility. Compared to conventional systemic cytokine administration or passive-release biomaterials, our approach offers mechanistic advantages, including localized and sustained cytokine presentation, active immune cell recruitment, and minimally invasive delivery. Collectively, these features position the hydrogel as a translationally relevant platform for kidney-targeted immunotherapy, with potential applications in AKI, prevention of CKD progression, and broader inflammatory renal disorders.

## 6. Description of Supporting Information Material

Supporting Information is available online free of charge (PDF).

## 7. Acknowledgements

This work was supported by Startup Funds from the School of Biomedical Engineering and Sciences at Drexel University, NIH R35GM147184, and the Penn Center for Musculoskeletal Disorders Histology Core (P30 AR069619).

## Notes

### Competing Interest Statement

The authors have declared no competing interest.

