## Supporting Information for "Avidity-controlled biotherapeutic delivery systems for the treatment of ischemia-reperfusion acute kidney injury"

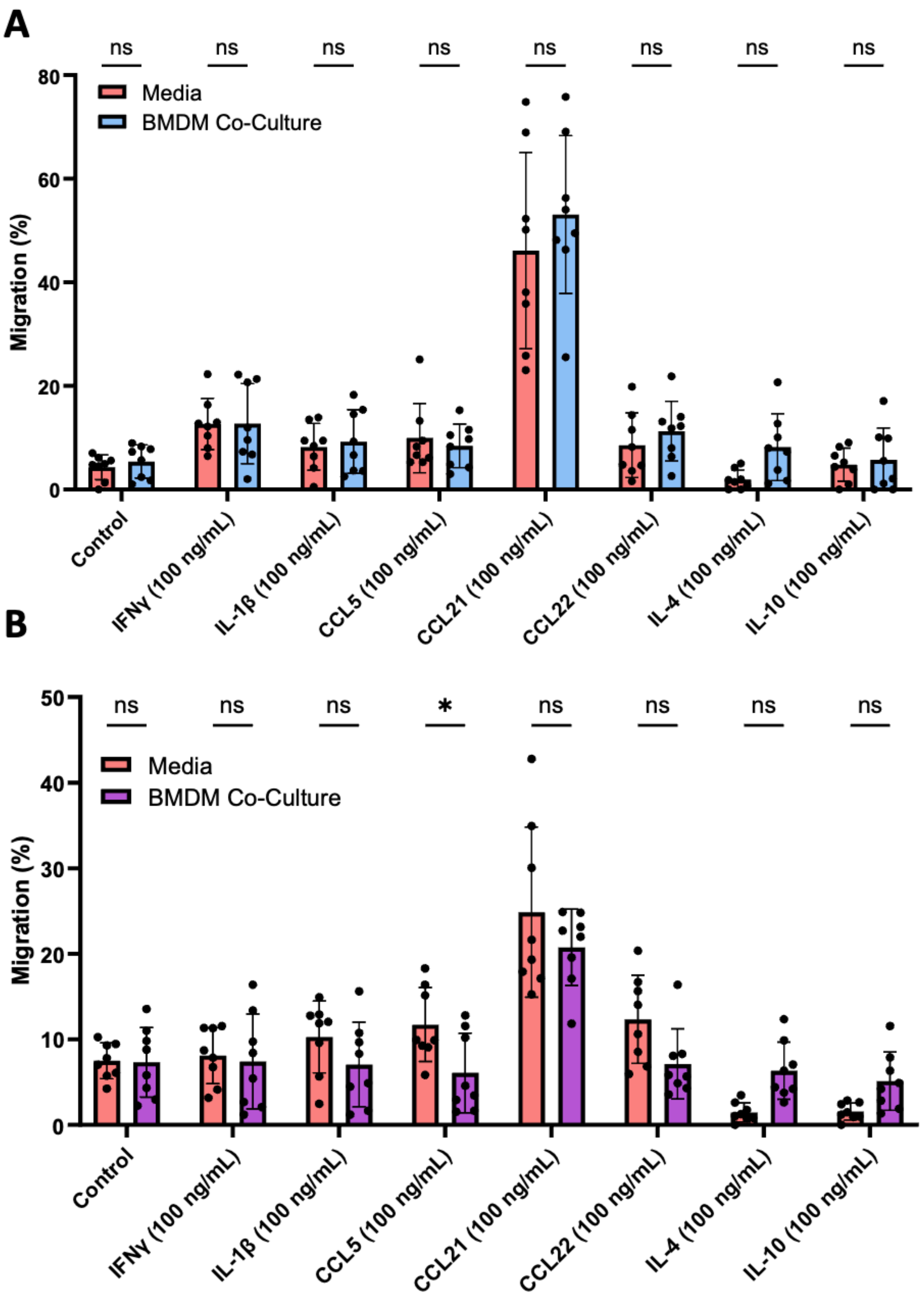


**Supplementary Figure 1. Transwell assays assessing T cell chemotaxis toward different signals.** Transwell migration assays of **(A)** CD4+ T cells and **(B)** Tregs toward BMDMs treated with chemoattractants compared to chemokine alone after 24 hours *in vitro*. Data are pooled from two independent experiments (n = 4 per experiment; total n = 8); *p<0.05; one-way ANOVA, Tukey HSD.

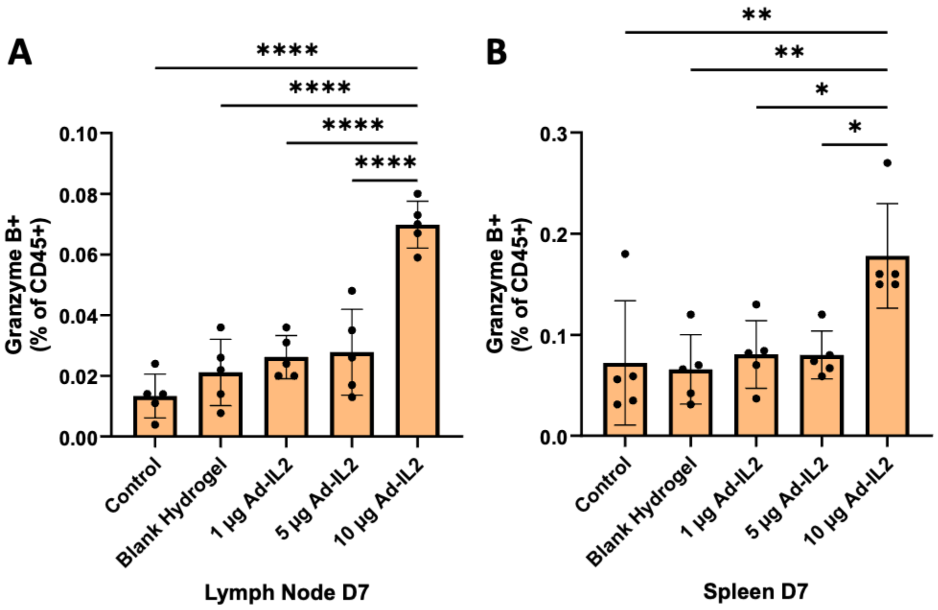


**Supplementary Figure 2. Assessment of cytotoxic T cells in lymphoid tissues after hydrogel injection in healthy mice.** Percentage of granzyme B+ T cells in the **(A)** lymph node and **(B)** spleen on day 7. Data represent mean ± SD, n=5; *p<0.05, **p<0.01, ****p<0.0001; one-way ANOVA, Tukey HSD.

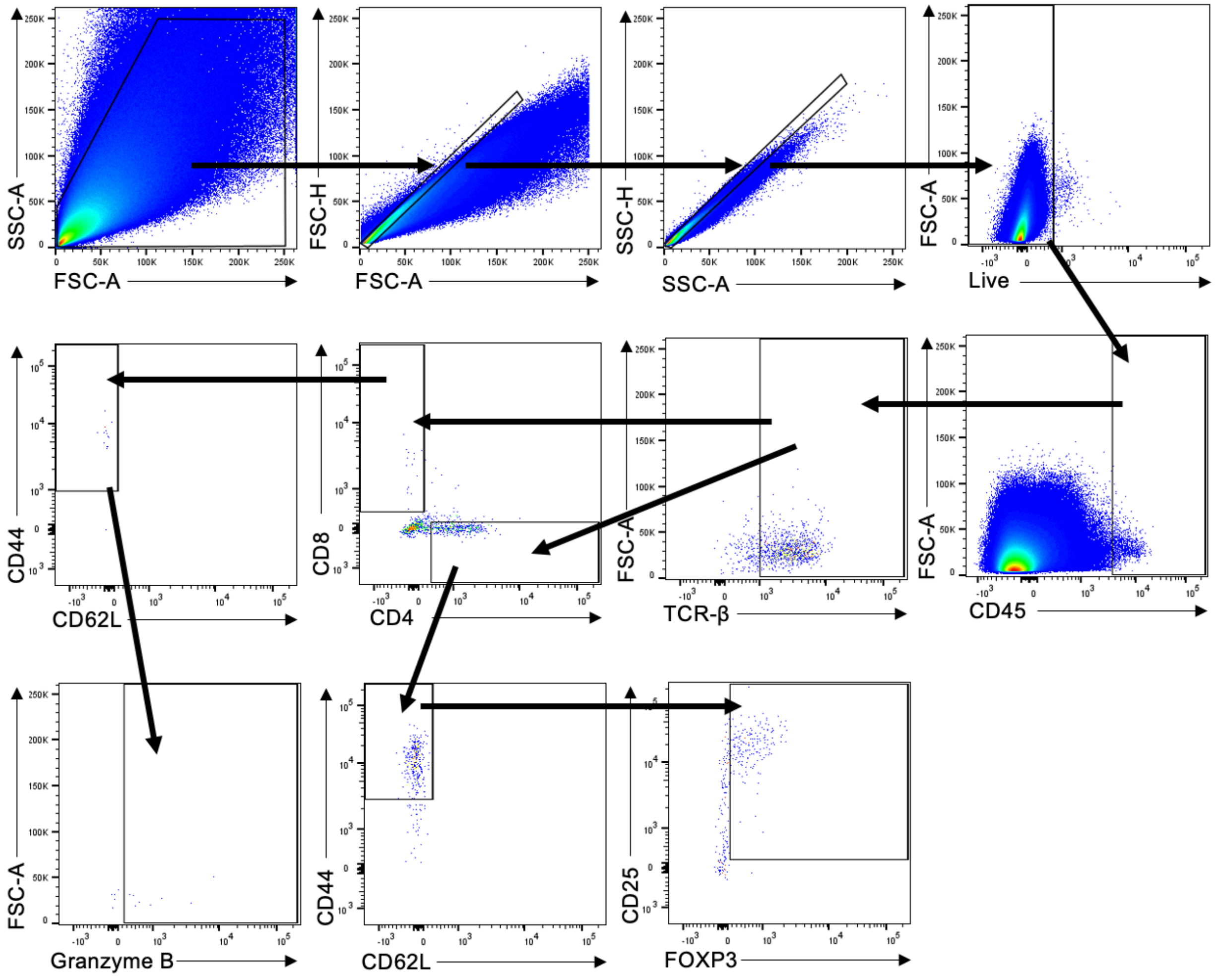


**Supplementary Figure 3.** Representative flow cytometry gating strategy for analysis of immune cells from subcutaneous hydrogel explants in healthy mice after 28 days.

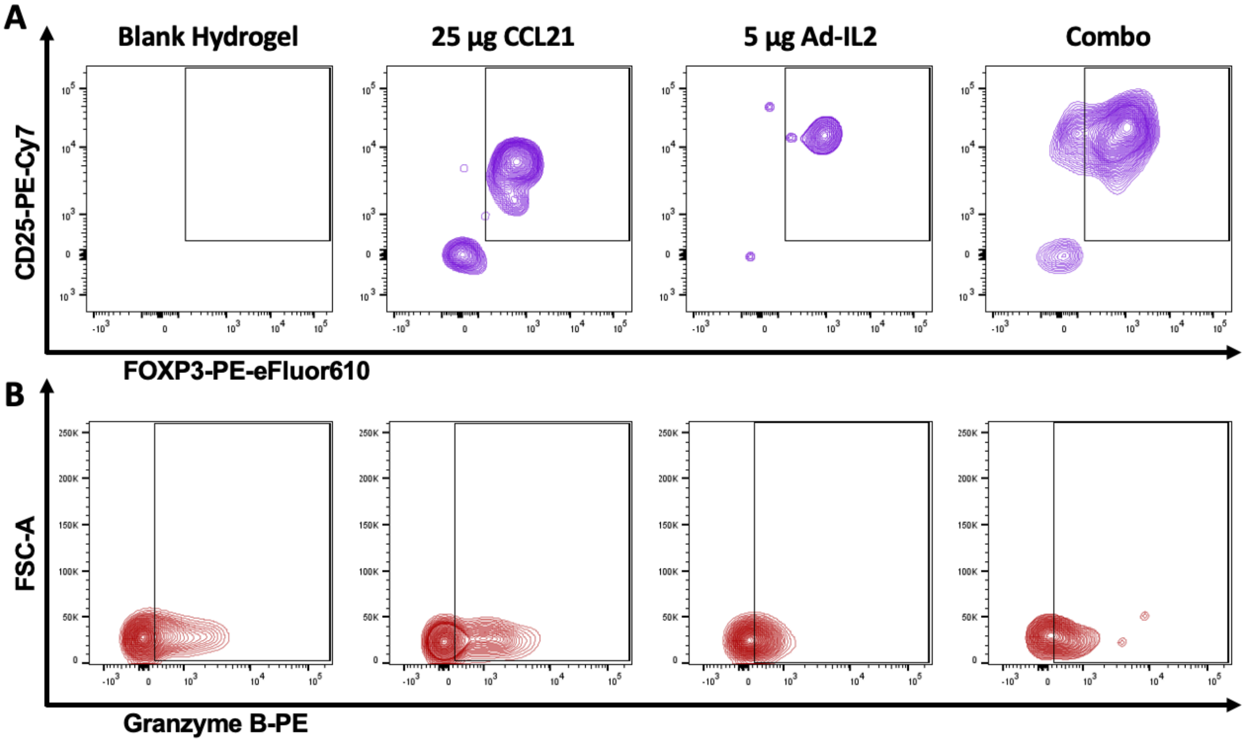


**Supplementary Figure 4.** Representative flow cytometry gating strategy for **(A)** Tregs and **(B)** **g**ranzyme B+ T cells in healthy mice at day 28 following subcutaneous hydrogel injection with blank, 25 µg CCL21, 5 µg Ad-IL2, and 25 µg CCL21 + 5 µg Ad-IL2.

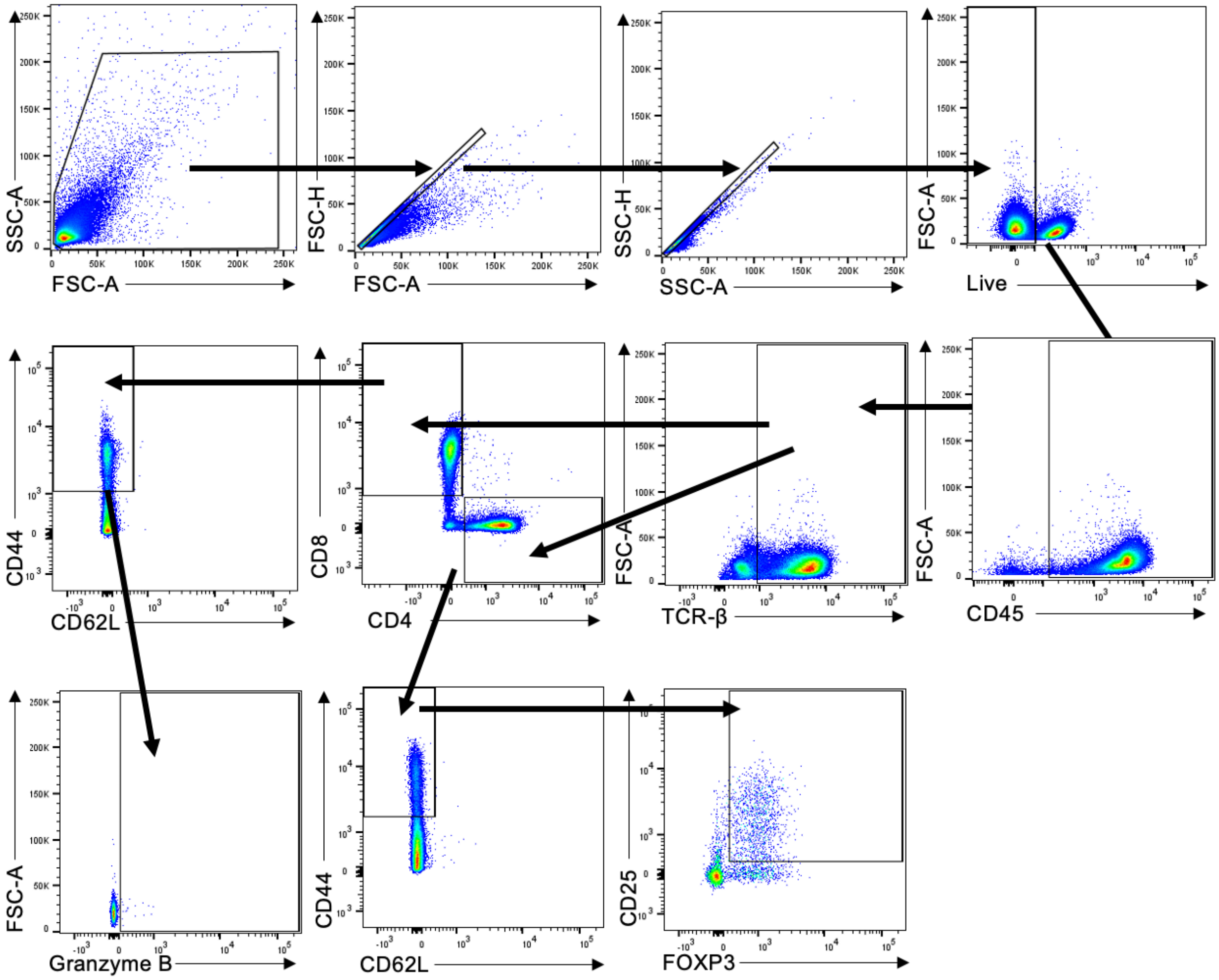


**Supplementary Figure 5.** Flow cytometry gating strategy for analysis of immune cells from spleen and lymph nodes of healthy mice 28 days after subcutaneous hydrogel injection.

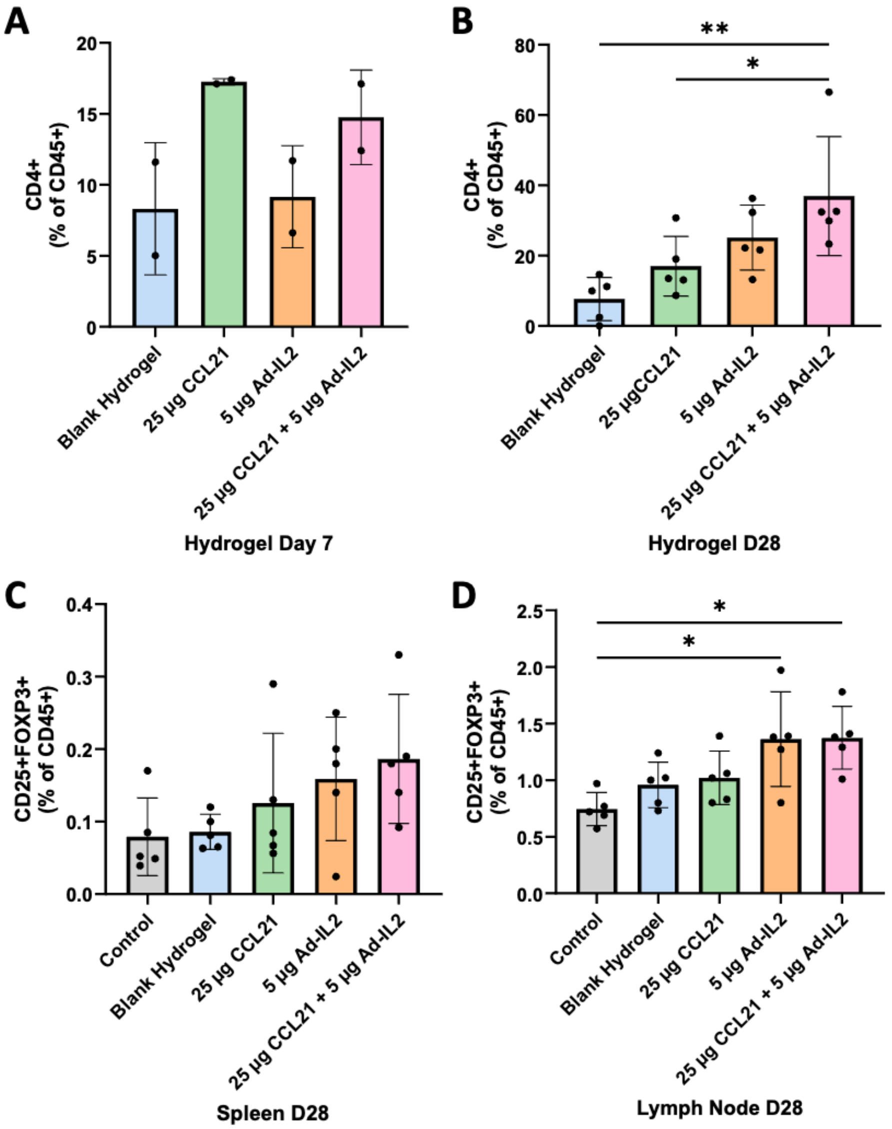


**Supplementary Figure 6. Immune cell composition in hydrogel explants and lymphoid organs in healthy mice.** Percentage of CD4+ T cells on **(A)** day 7 and **(B)** day 28 in hydrogel explants. Percentage of Tregs in the **(C)** spleen and **(D)** lymph node on day 28. Data represent mean ± SD, n=2 (A), n=5 (B-D); *p<0.05, **p<0.01; one-way ANOVA, Tukey HSD.

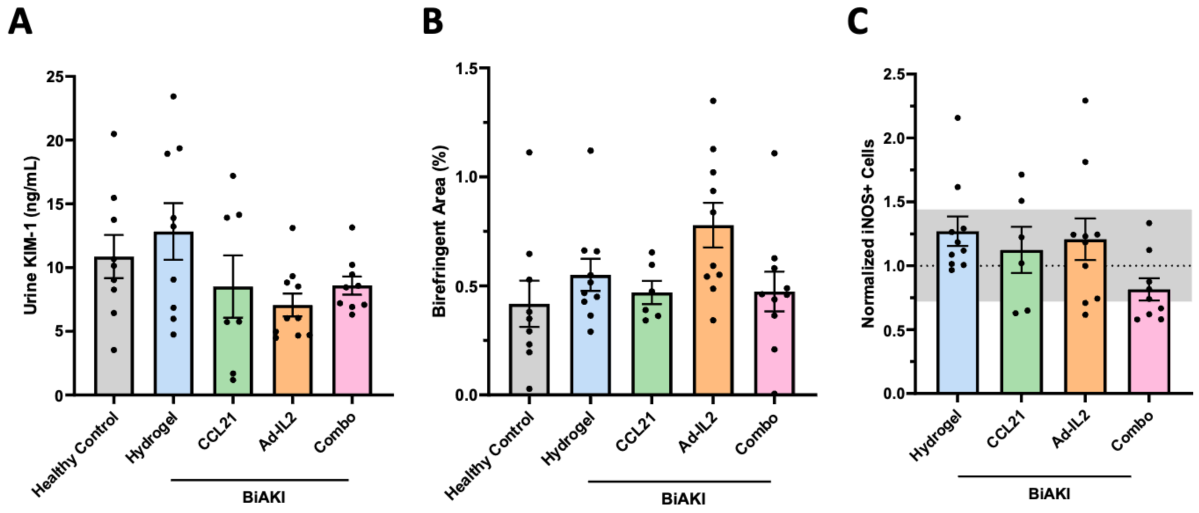


**Supplementary Figure 7. Assessment of fibrosis, kidney injury, and inflammatory markers at day 28 post-BiAKI induction. (A)** Urinary kidney injury molecule-1 (KIM-1) levels measured at day 28. **(B)** Quantification of percent birefringent area in the outer cortical rim of picrosirius red (PSR)-stained kidney sections imaged under polarized light, indicating collagen deposition and fibrosis. **(C)** Quantification of normalized inducible nitric oxide synthase (iNOS) cells from immunofluorescence analysis. Data represent mean ± SEM, n ≥ 6.

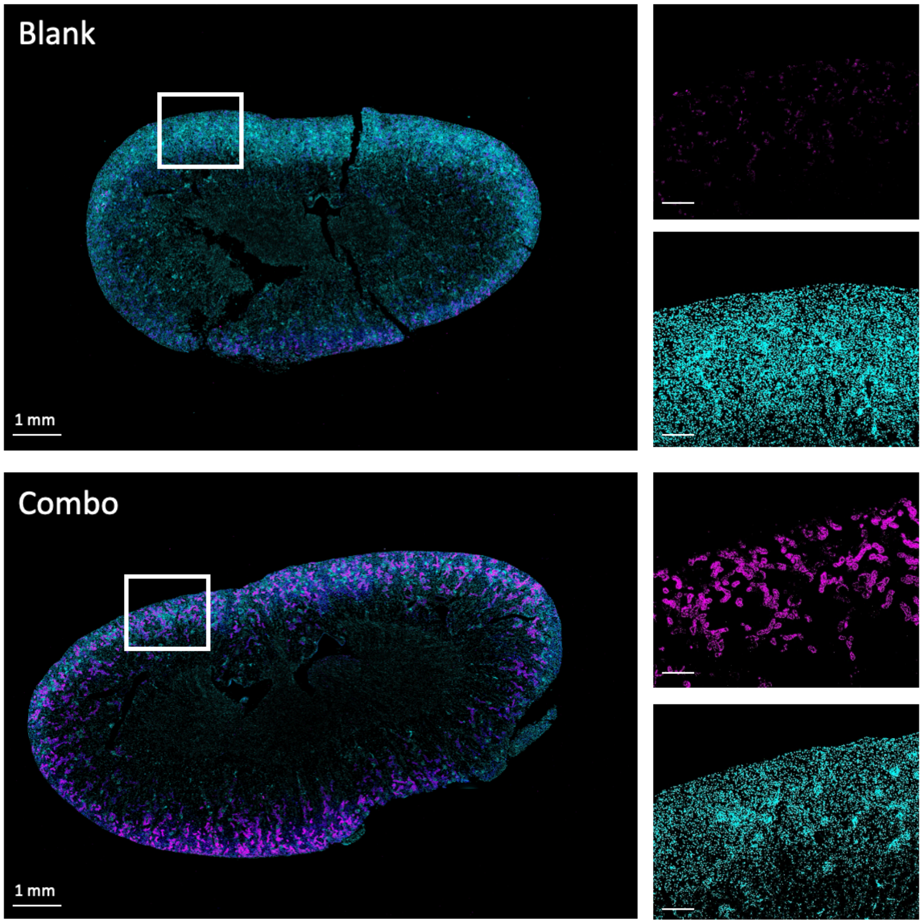


**Supplementary Figure 8. Whole-kidney overview and cortical immunofluorescence at day 28 post-BiAKI induction**. Representative images from kidneys treated with blank hydrogel and CCL21 + Ad-IL2–loaded combination hydrogel. Whole-kidney overview images show merged DAPI and FOXP3 staining, followed by higher-magnification views of the cortex with corresponding individual channels (DAPI and FOXP3). Images confirm cortical localization used for quantification. Scale bar 250 μm unless otherwise indicated.

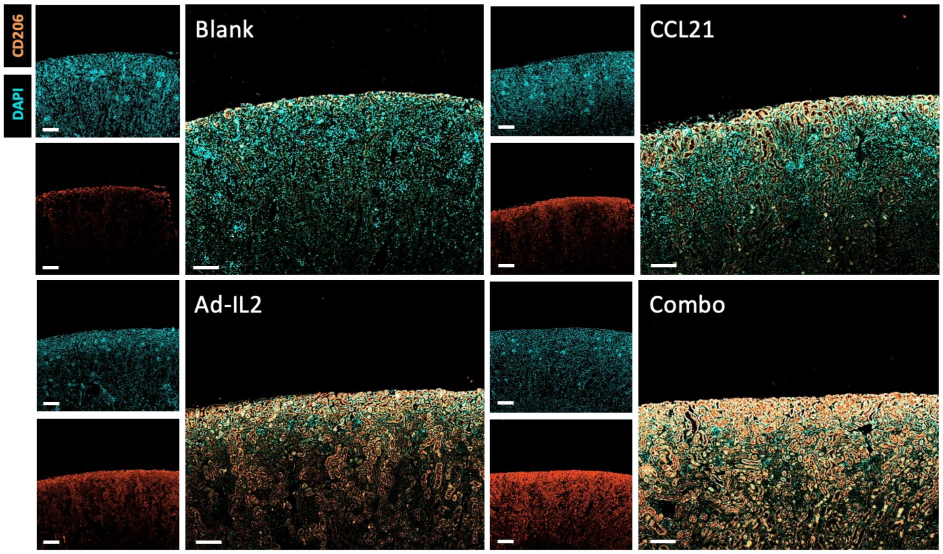


**Supplementary Figure 9. CD206⁺ macrophage localization in the renal cortex following hydrogel treatment after BiAKI**. Representative immunofluorescence images of BiAKI kidneys treated with blank, CCL21, Ad-IL2, or combination hydrogel. Images show merged DAPI and CD206 staining from the renal cortex, with corresponding single-channel views (DAPI and CD206) from the same field of view. Scale bar 250 μm.
